# CLOOSE: An open-source platform for optical closed-loop experiments

**DOI:** 10.1101/2025.04.09.648061

**Authors:** Valerio Francioni, Anna Beltramini, Linlin Z Fan, Mark T. Harnett

## Abstract

Brain-Computer Interfaces (BCI) have catalyzed advancements in both clinical applications and basic neuroscience research. However, technical barriers such as steep learning curves and complex synchronization requirements often impede their widespread adoption. In response to the increasing demand for optical BCI experiments and to the technical challenges associated with their implementation, we introduce CLOOSE (Closed-loop Optical Open-Source Experiments), a versatile platform designed to standardize, simplify, and accelerate closed-loop experiments with functional imaging data. CLOOSE interfaces easily with any frame-based imaging system via TCP/IP, allowing for real-time data streaming and distributed computation. Benchmark tests validate CLOOSE’s real-time accuracy in image registration, signal processing, and analysis at imaging frequencies ≥1 kHz, making it the first optical BCI compatible with voltage indicators. Throughout the paper we showcase CLOOSE’s versatility in supporting several neurofeedback paradigms: from single neuron to population dynamics, multiplane imaging, and online (and offline) z-tracking. CLOOSE’s functionality is easily accessible to users with minimal coding experience through an intuitive graphical interface for experiment setup and real-time performance monitoring. By significantly lowering the barrier to performing technically demanding closed-loop experiments, CLOOSE advances the streamlining, standardization, and democratization of online image analysis and neurofeedback BCI paradigms, opening new avenues for precise manipulation and real-time readout of neuronal activity in health and disease.

## Introduction

Recent years have witnessed a remarkable increase in the use of BCI systems^1–3^. By providing direct and often bi-directional communication between real-time brain activity and external devices, BCIs have proved to be particularly useful for clinical applications requiring online analysis of brain activity such as neuroprosthetics^4–7^, speech synthesizers^8–11^, but also treatments for conditions such as chronic pain^12^, epilepsy^13,14^, Parkinson’s disease^15,16^, depression^17^, and sleep disorders^18,19^, among others^2^. In addition to clinical applications, BCIs have emerged as powerful tools for basic research. By allowing experimenters to determine the mapping across neuronal activity, external feedback stimuli, and reward, BCI systems provide a highly controlled approach to studying learning, synaptic plasticity, motor control, population dynamics and credit assignment in vivo^20–30^. Additionally, unlike traditional open-loop systems, closed-loop systems enable the experimenter to manipulate neuronal activity in a functionally and temporally precise manner, thereby facilitating the dissection of the circuit elements required for neuronal computations^31–34^. Combined with the advent of genetically-targeted opsins and novel methods for spatially-precise light delivery, the online manipulation of functionally-defined neuronal circuits has the potential to revolutionize the study of brain processes in health and disease^35^.

Despite the many advantages offered by closed-loop systems, their widespread adoption in neuroscience has been hindered by several challenges^2^. One significant obstacle is the steep learning curve associated with implementing and operating experiments requiring online analysis of large neuronal data acquired at high speeds. Designing these experiments requires specialized expertise in areas such as software engineering, online signal processing, experimental design, and data analysis, making these tools inaccessible to many researchers without extensive training. Moreover, the technical complexity involved in synchronizing multiple components, including neuronal recording devices, feedback delivery systems, reward delivery, and other accessory external devices (such as LEDs for optogenetics) further increases the barrier to entry for conducting closed-loop experiments. Due to these complexities, closed-loop firmware is often implemented to work exclusively within the boundaries of one’s experimental setup, often behind the curtain of proprietary software^36^. As such, closed-loop experiments are hard to customize and are inaccessible to the wider community, and the interoperability and standardization is encumbered by the lack of a common platform.

Here, we introduce CLOOSE (Closed-Loop Optical Open-Source Experiments), a versatile MATLAB-based platform for conducting closed-loop optical experiments. CLOOSE provides multiple avenues for short-latency online data analysis while also supporting offline processing options. CLOOSE interfaces with any frame-based imaging acquisition devices via a standard Transmission Control Protocol/Internet Protocol (TCP/IP) connection. TCP/IPs provide fast and reliable data transfer over multiple network types including Local Area Networks (LANs), Wide Area Network (WANs), and Peer-To-Peer (P2P) networks with wide compatibility across device types, regardless of hardware and operating system. As such, CLOOSE can be run on a separate machine as the main data acquisition PC, allowing the experimenter to distribute the computational load across multiple PCs for faster performance. Its graphical user-friendly interface (GUI), modular software design, and open-source code facilitate integration with various acquisition systems and experimental setups, thus standardizing, democratizing, and decentralizing access to closed-loop methodologies within the wider neuroscience community. We showcase the speed and versatility of CLOOSE in several experimental contexts underscoring its potential for advancing the understanding of brain function and behavior. Additionally, we provide comprehensive documentation and code examples to facilitate the adoption and customization of CLOOSE for specific research needs, thereby fostering a collaborative and open-source approach to closed-loop optical experiments. All CLOOSE code is available at the GitHub (https://github.com/harnett/CLOOSE) repository together with extensive documentation, examples and demos.

## Results

### CLOOSE workflow and performance

CLOOSE is an open-source, standalone software package written in MATLAB R2024b that flexibly interfaces with any image acquisition system to perform a variety of closed-loop experiments and online analyses. CLOOSE operates on serially streamed data from an acquisition PC via a TCP/IP connection. Compared to other protocols such as User Datagram Protocol (UDP), TCP/IP connections are more reliable and scalable, and compatible with a wide range of applications. CLOOSE can be installed either on the same or a different machine as the data acquisition PC (Figure 1a). This gives the experimenter the option to distribute computational and data loads across multiple devices for maximized performance and speed. Individual 2D frames (.bin or .tiff) of any arbitrary user-defined dimensions (x and y pixels) and quality (e.g., uint8 or uint16) are streamed to CLOOSE as 1D image vectors (Figure 1b-d). While we focus on neuronal data acquired using either one- or two-photon microscopes, CLOOSE only requires a serial stream of imaging data, and it can therefore seamlessly integrate with virtually any image acquisition device. The core processing of individual 2D frames received by CLOOSE consists of five steps (Figure 1e): **1) Image processing -** the 2D reconstruction of individual frames received as 1D vectors, according to user-defined parameters. **2) Motion correction (or image registration, optional) -** online image registration is crucial to perform accurate close-loop experiments, as it mitigates the effects of subject movement, enhancing the quality of data for further analysis and accurate mapping between neuronal activity and the external device. CLOOSE performs the rigid re-alignment of a full frame, based on n_q_ sub-frames selected from a reference image, where the area (NqX x NqY), location and total number of subframes (n_q_) are all user-defined parameters (Figure 1g-i) which can be customized based on the experimental conditions (e.g., sparse vs dense labelling). Reducing either n_q_ or NqX/NqY will increase the speed of motion correction. The median x and y pixel displacement for all n_q_ quadrants are used to apply a global frame registration (Figure 1h). The reference image to which each new incoming frame will be aligned can either be acquired de novo using CLOOSE, or uploaded from a previously acquired session. See wiki (https://github.com/harnett/CLOOSE/wiki) for more details. The online image registration algorithm incorporated within CLOOSE is precise with subpixel resolution (see Methods)^37^. While retaining a latency of 1.6 ms per quadrant (128x200 pixels), when tested on real data (Figure 1f), CLOOSE differs from other commonly-used offline motion correction algorithms^38^ by 0.15 pixels (in both x and y displacement) per frame, on average (N = 30 imaging sessions, Figure 1i). **3) Fluorescence extraction -** the Regions of Interest (ROIs) are defined by the experimenter prior to the beginning of the experiment. ROIs can be either drawn online or uploaded from previous sessions. ROI drawing is based on a click-based free-shaped polygon tool, which allows for subsequent modifications (i.e., fine edge adjustments), displacements and deletions. This approach makes ROI-drawing amenable for diverse morphologies including cell bodies, spines, or other neurites. **4) Plotting (optional) -** live visual feedback consisting of newly acquired 2D frames, online-extracted fluorescent traces and performance tracking for the experimenter to assess the quality of their data in real time. **5) Feedback stimulus presentation (optional) -** CLOOSE provides the experimental subject with information about their performance during neurofeedback tasks. CLOOSE comes with the option to deliver either visual or auditory stimuli using Psychtoolbox^39^ and MATLAB internal audio player, respectively.

**Figure 1:**
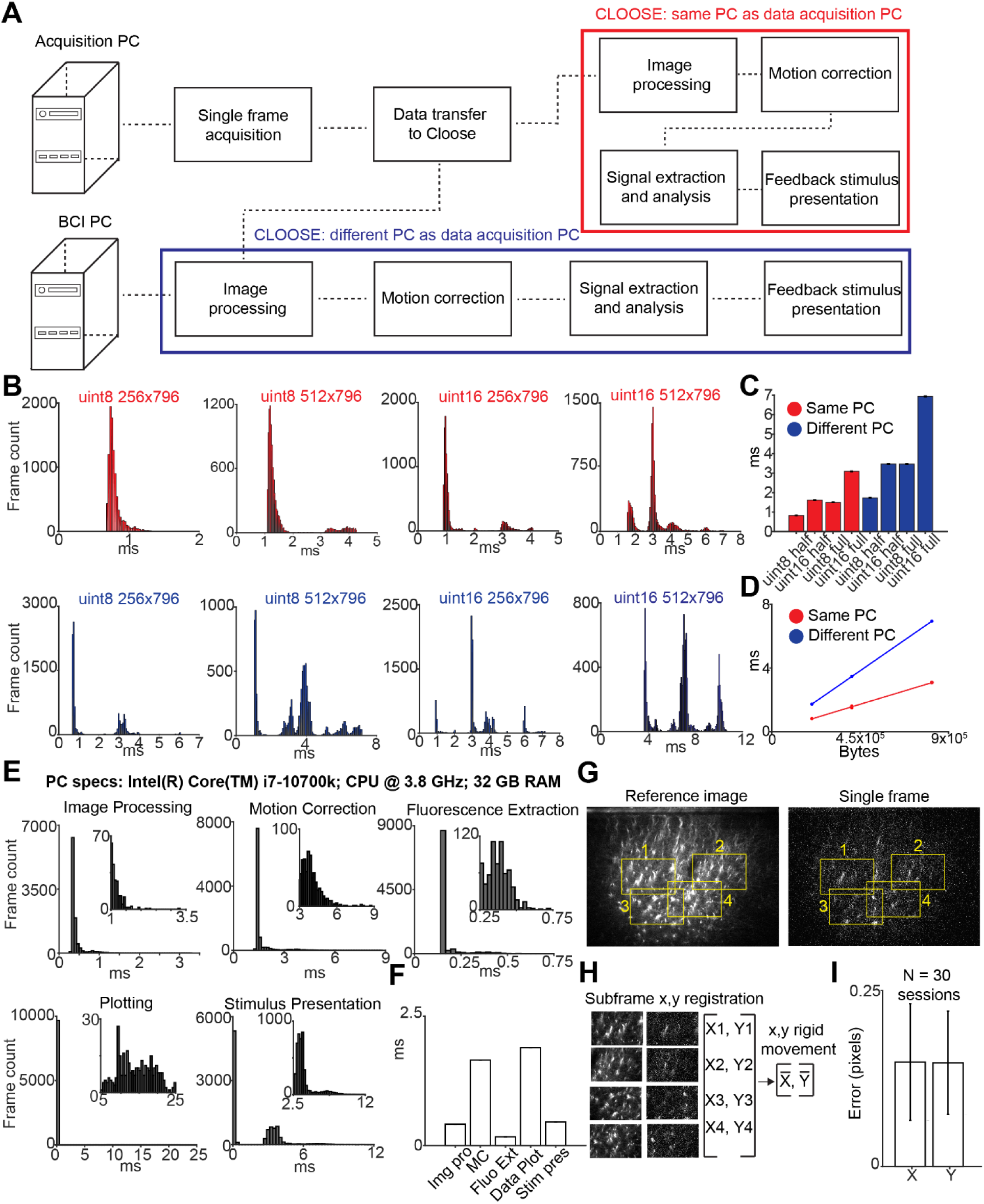
CLOOSE workflow and performance. A, Schematic representation of the different processing steps performed by CLOOSE. CLOOSE can be installed either on the same PC as the data acquisition PC (in red), or on a separate PC (in blue) which data are transferred to from the acquisition PC. **B,** Data transfer speeds for data of different quality and size. Each datapoint represents one new frame for a total of 10000 frames. Upper row (red) shows latencies for transfers between two software installed on the same PC. Lower row (blue) shows latencies for transfers on two different PCs. Uint8 and uint16 data have individual pixel values ranging from 0 to 255 (1 byte) and from 0 to 65535 (2 bytes), respectively. **C**, Mean transfer latencies for the histograms shown in b. We define a full image as a 512x796 pixels image and half as a 256x796 pixels image. **D**, Mean transfer latencies for the histograms shown in b as a function of total bytes transferred. **E**, For each of the processes performed by CLOOSE, the histogram of the latencies for various processing steps. Each datapoint represents one new frame. The steps were assessed on real data processed using a PC whose specifications are reported on top of the panel. **F**, Mean transfer latencies for the histograms shown in e. For motion correction, the value represents the latency to align one subframes each of 199x128 pixels at uint16 quality. **G,H**, A step-by-step breakdown of motion correction, in this case using four subframes each of 199x128 pixels at uint16 quality. Each newly-acquired single frame is decomposed into four subframes at some user-defined location. These subframes are compared against the subframes, located in the same location, of a previously acquired reference image. Using a Fourier-transform approach (see methods), an x and y pixels shift is estimated for each subframe, the median of which is used to apply a global rigid 2D image translation. **I**, Mean absolute error, defined as the average difference between the offline motion correction algorithm of Suite2P and the motion correction approach used by CLOOSE. Throughout the figure, error bars represent standard error of the mean (SEM).

CLOOSE demonstrates effective handling of large-scale neuronal datasets with high accuracy and speed: CLOOSE can perform a full processing loop in under 5ms on a low-to-medium tier PC (Figure 1e). In Figure 5, we demonstrate how this performance can be further enhanced for applications using super-fast reporters including voltage indicators acquired at 1 kHz.

### CLOOSE supports diverse neurofeedback experimental designs: from single cell to population dynamics

Different hypotheses require distinct experimental designs. However, bottom-up adaptation of software architecture can be cumbersome and often introduces hard-to-detect bugs and performance bottlenecks due to conflicts with higher level dependencies. To address this challenge, we optimized CLOOSE to seamlessly switch between three commonly-used experimental designs in neurofeedback BCI tasks. Across all three tasks, neuronal activity is mapped to a feedback stimulus (visual or auditory) which reports how close (in terms of neuronal activity) the subject is to target – the neural activation required to trigger a secondary event such as a reward or the activation of an optogenetic probe.

**1) Single neuron/population design** - under this task design, both the feedback stimulus and the target are controlled by the activity of a single neuron or a single population of neurons (Figure 2a-c)^40^. A use-case for this approach is to investigate how systematic pairing of synaptic activity with rewards might lead to changes in spine activity or morphology.
**2) Two population design -** under this experimental regime, the feedback stimulus and the target are operated by the difference in activity between two neurons, two spines, or two populations of neurons (Figure 2d-f). This approach has been successfully utilized to study how two populations diverge as a function of their pairing - or anti-paring - to a reward ^25,26^ and how instructive signals differ in these two populations – a potential approach to address biological credit assignment^24^.
**3) The network dynamics approach -** under this experimental regime, the feedback stimulus and the target are operated when neuronal population activity occupies a specific point within an n-dimensional space (Figure 2g-i). This approach, has been utilized to reinforce physiological neuronal activity patterns^29,41^. CLOOSE supports analysis in n-dimensional space, where n equals the number of neurons selected by the user. Alternatively, CLOOSE can project vectors of neuronal activity onto dimensionally-reduced spaces such as Principal Components or t-distributed Stochastic Neighbor Embeddings (tSNE) in n_c_ dimensional space (Figure 2g-i) where n_c_ is user-defined in the case of principal components and equals 2 for tSNE.

**Figure 2:**
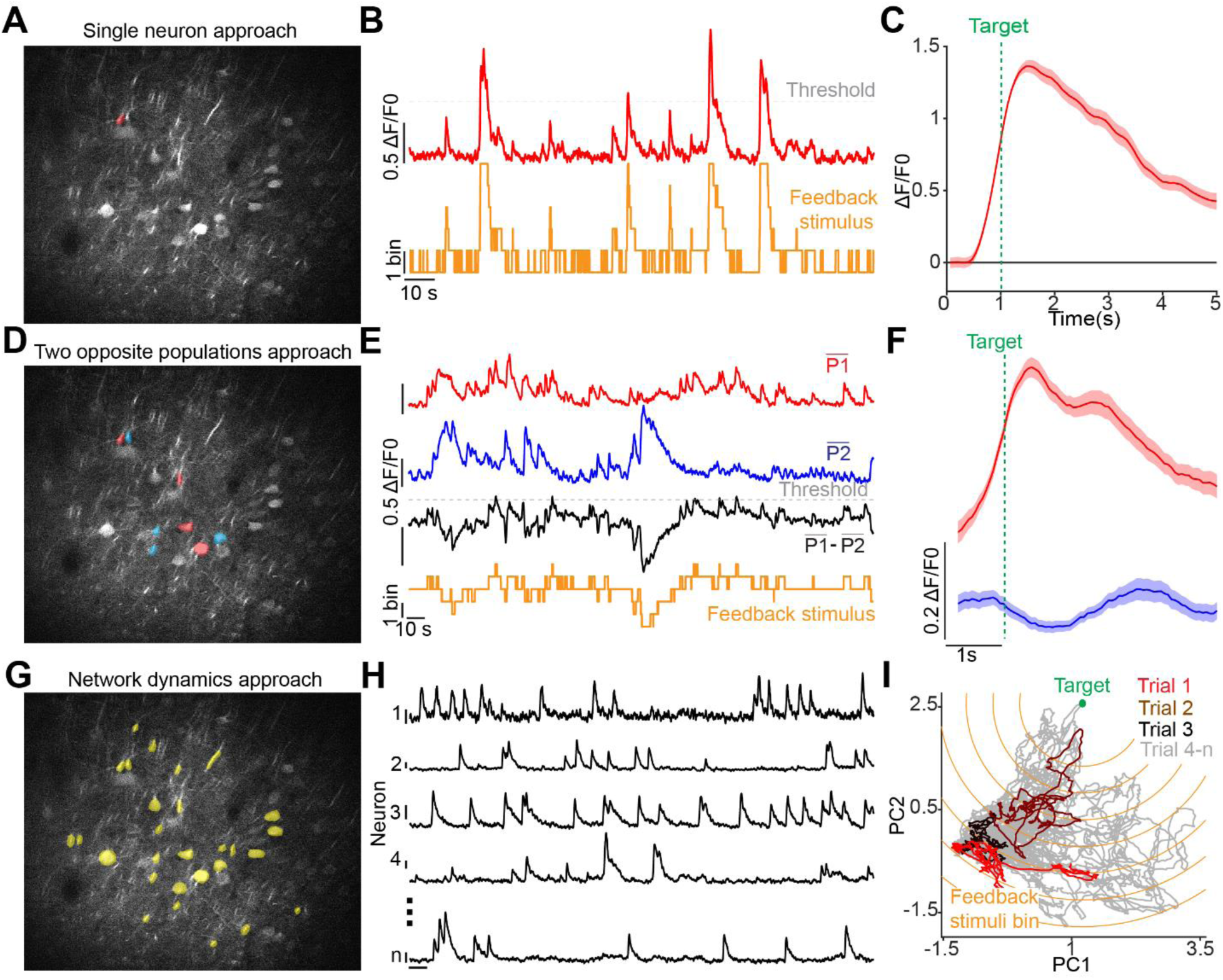
CLOOSE supports a range of different experimental designs: from single cell to population dynamics. **A**, Field of view of GCaMP7f-labelled layer 5 neurons with a ROI (in red) over the cross-section of the apical trunk of a single neuron to exemplify the single neuron approach to neurofeedback experiments. **B**, The activity of a single neuron quantified as changes in fluorescence (ΔF/F0) is alone responsible for driving the feedback stimulus (segmented in 7 bins). ΔF/F0 and feedback stimulus are linearly mapped to one another. The grey dashed lines indicate the activity threshold to trigger an output (e.g. reward or an optogenetic stimulus). **C**, The mean ΔF/F0 at target (from one second before to four seconds after target reach). **D**, Same FoV as shown in a, for a two opposite populations task design. Red, neurons whose activity drives the feedback stimulus towards a target, (P1). Blue, neurons whose activity drives the feedback stimulus away from the target (P2). **E**, The mean ΔF/F0 for the P1 and P2 populations is estimated and subtracted from one another (black trace). This subtracted activity is linearly mapped to a feedback stimulus during the closed loop part of a neurofeedback BCI task. **F**, Same as c, mean activity for P1 and P2 neurons at target. **G**, Same FoV as in a and d for a network dynamics task design with ROIs highlighted in yellow. **H**, Activity of five representative neurons among the ones highlighted in g. **I**, At each time point, the n-dimensional activity of the neurons selected in a is collapsed onto the first two principal components. The Euclidean distance from a target network state is estimated in a moment-by-moment manner and the feedback stimulus is delivered according to a system of concentric circles around the target. Three individual closed-loop trials (in color) are highlighted over all the other trials (in grey) of a single training session.

Across all three different approaches, experimenters can select parameters, including the identity of the ROIs partaking in the task, the fraction of trials in which neuronal activity reaches the target (effectively setting the difficulty of the task), as well as the number of bins in which the feedback stimuli are subdivided.

### CLOOSE supports volumetric BCI

Different hypotheses also require BCI software to be compatible with diverse imaging approaches. Recent advances in piezoelectric systems or electrically-tunable lenses allow fast scanning of multiple 2D planes to generate a 3D stack and maximize the number of individual units recorded. CLOOSE is equipped to handle volumetric data in two separate ways: **1) Multiplane BCI -** CLOOSE is designed to allow the experimenter to run BCI experiments using ROIs drawn on multiple independent planes. Using this approach with imaging in animal models, experimenters could utilize the network dynamics of an entire cortical column or the joint activity of somas and dendrites to drive the BCI (Figure 3a, b). **2) Single plane volumetric BCI** - users can run the BCI using activity from one or more imaging planes while simultaneously recording from other imaging planes which do not have a causal role in driving the BCI (Figure 3c). Our group used this approach in mice to separately record dendritic activity of layer 5 neurons while their corresponding somas were used to run the BCI task^24^. Alternatively, this approach can be used to study how the activity of a specific layer evolves as a function of the changes induced by the neurofeedback task in a different cortical layer (Figure 3d).

**Figure 3:**
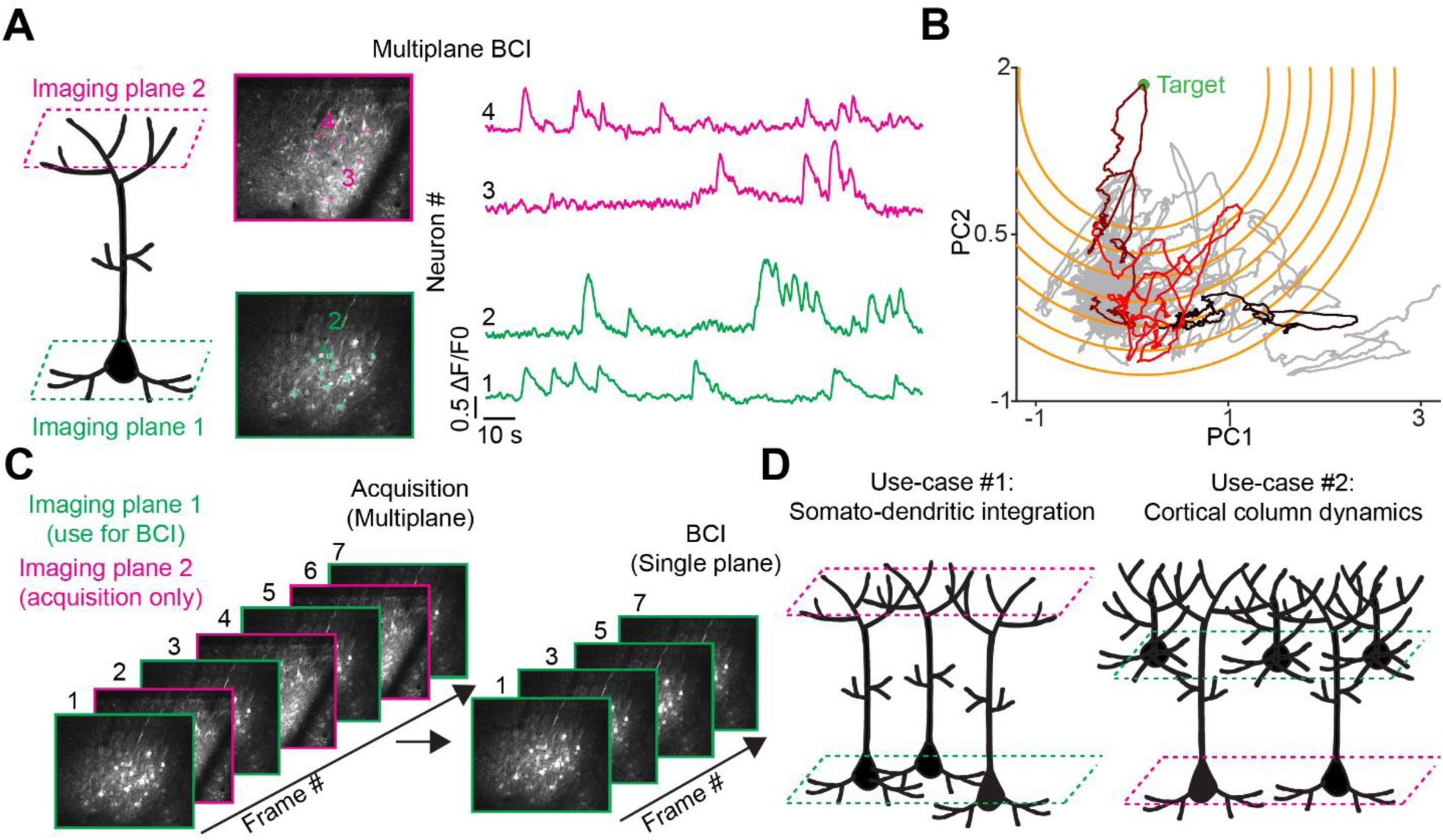
CLOOSE supports volumetric BCI. A, Left Panel, a schematic of the imaging approach. Middle, two FoVs, recorded 400 (green) and 150 (purple) um below the dura depicting the soma and dendrites of layer 5 pyramidal neurons labelled with GCaMP7f. Ten ROIs per plane are used to run the BCI using the network dynamics experimental design. ΔF/F_0_ traces for four (two per plane, in green and purple) neurons acquired simultaneously (15 Hz per plane) using an ETL. **B**, At each time point, the 20th-dimensional activity of the neurons selected in a is collapsed onto the first two principal components to estimate the Euclidean distance from target activity and its corresponding feedback stimulus. Three individual closed-loop trials (in color) are highlighted over all the other trials (in grey) from a single training session. **C**, Same acquisition settings as in a but with only the lower plane (somas, in green) used to run the BCI. CLOOSE allows the user to run BCI experiments while simultaneously acquiring data from other imaging planes without interference. **D**, Schematics for two hypothetical use-cases for the experimental design illustrated in c.

### Online (and offline) z-tracking using CLOOSE

CLOOSE’s capacity for managing volumetric data is further demonstrated by its z-tracking feature (Figure 4 a-d). During imaging, both abrupt z-shifts as well as slow z-drifts lead to blurred and distorted frames that introduce signal artifacts, compromising data quality and the subsequent analysis. Importantly, these artifacts cannot be corrected offline. As a consequence, identifying (online and offline) and correcting (online) for these drifts is crucial to ensure interpretable results. Users can exploit the z-tracking functionality in CLOOSE to online - or offline (“alignment only” checkbox in the GUI) -track motion in the z-dimension. When used during chronic imaging experiments, this functionality maximizes the number of neurons that can be tracked across days by enabling the experimenter to fine-tune the focal plane to match fields of view over time. Similar to the image registration function, users can either acquire a volumetric z-stack de novo, prior to running their experiment, or upload a previously acquired z-stack of reference images. Reference frames composing the 3D stack can then be used for comparison against each newly-acquired, motion-corrected frame. To test the algorithm’s precision on ground-truth data, we imaged pollen grains (a static reference), at multiple depths separated by 1 μm and compared each newly-acquired frame against a z-stack of reference images spanning a total of 20 μm (21 frames 1 μm apart) (Figure 4b). For each newly-acquired frame, we computed the mean squared error (MSE) value against each of the 21 reference frames (Figure 4c). The best match was selected to be the one with the smallest MSE. Our results demonstrate the high degree of accuracy in matching over 90% of all the individual frames within 1 μm from ground truth at high speed – below 2 ms for a comparison against a z-stack of 21 images (Figure 4c, d). Notice that for errors of 1 to 2 um where the vast majority of the distribution lies, the difference is hardly perceptible also for a trained human eye. Experimenters can customize multiple aspects of this functionality including the distances between two consecutive planes in the reference z-stack, the size of the area used to estimate the MSE, or by reducing the total number of planes within a given stack to compare each frame against. These optimizations reduce the number of total comparisons and increase processing speed and accuracy based on the user’s requirements. Users might also utilize the output of this function to adjust their focus online (e.g., using a piezoelectric motor or an Electrically-Tunable Lens (ETL)) or to correct a time-averaged version of slow z-drifts by adjusting the stage position. This functionality is ideally suited for imaging and performing closed-loop experiments on small structures such as dendrites and spines, where changes in the z-plane can be the most disruptive for accurate data collection.

**Figure 4:**
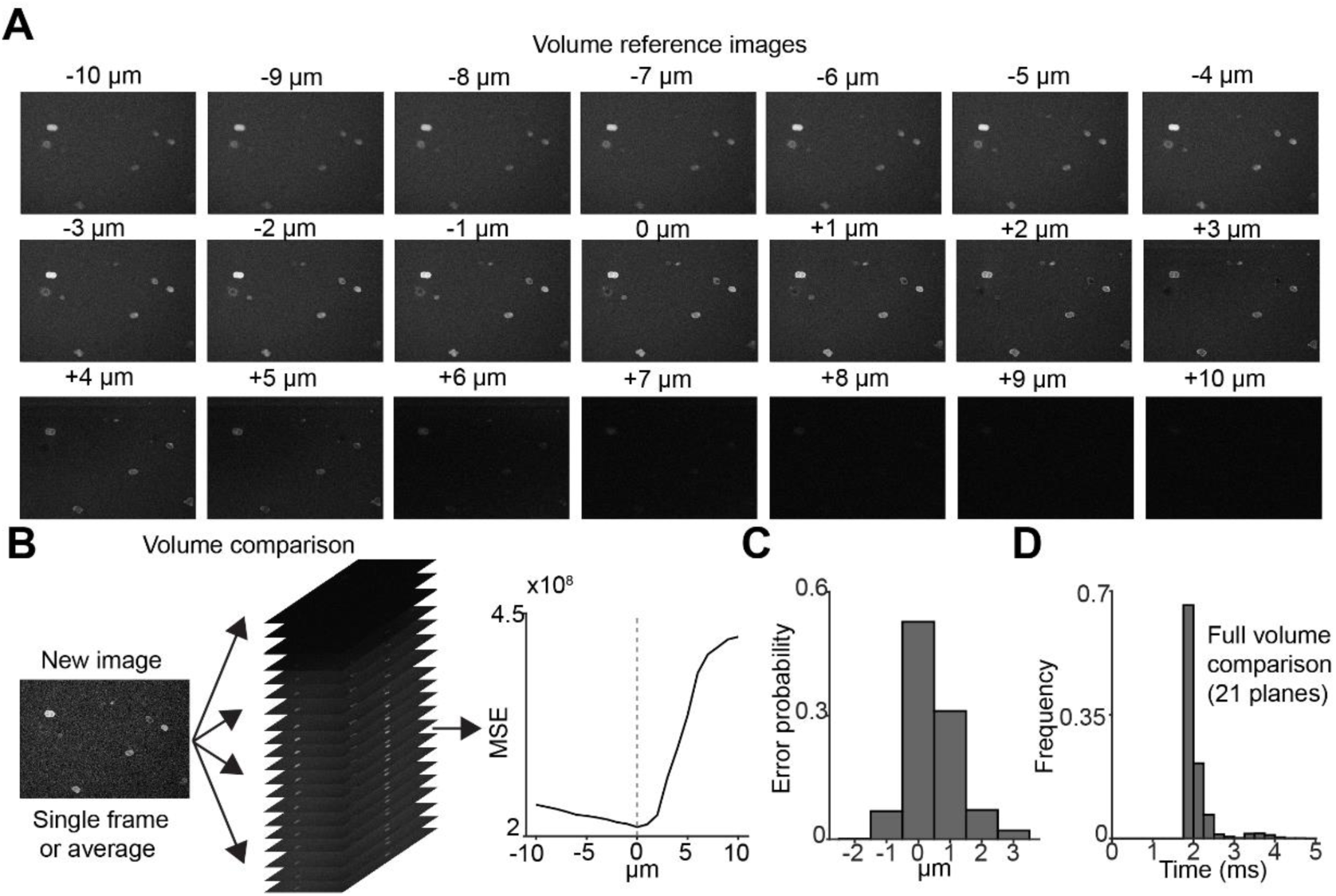
Online (and offline) z-tracking using CLOOSE. A, Volume reference images (50 frames average each) spanning 20 um along the z-axis acquired using an ETL. Each of the images is 1 um apart from the next. **B**, Each newly acquired image is first motion corrected against its reference image (not shown) and after that, is compared against each individual reference z-stack (21 individual comparisons, left panel) and the mean squared error is estimated for each 2D reference in a 3D stack (right panel). The one plane where the MSE is the lowest is the estimated z depth of the newly-acquired data. **C**, Error distribution compared to ground truth. Notice that for errors of 1 to 2 um where most of the distribution lies, the difference is hardly perceptible also for a trained human eye. **D**, For each image, the latency (in ms) for online estimating the MSE of that image against all the 21 reference images. Note that processing can be sped up by comparing the image against smaller volumes.

**Figure 5:**
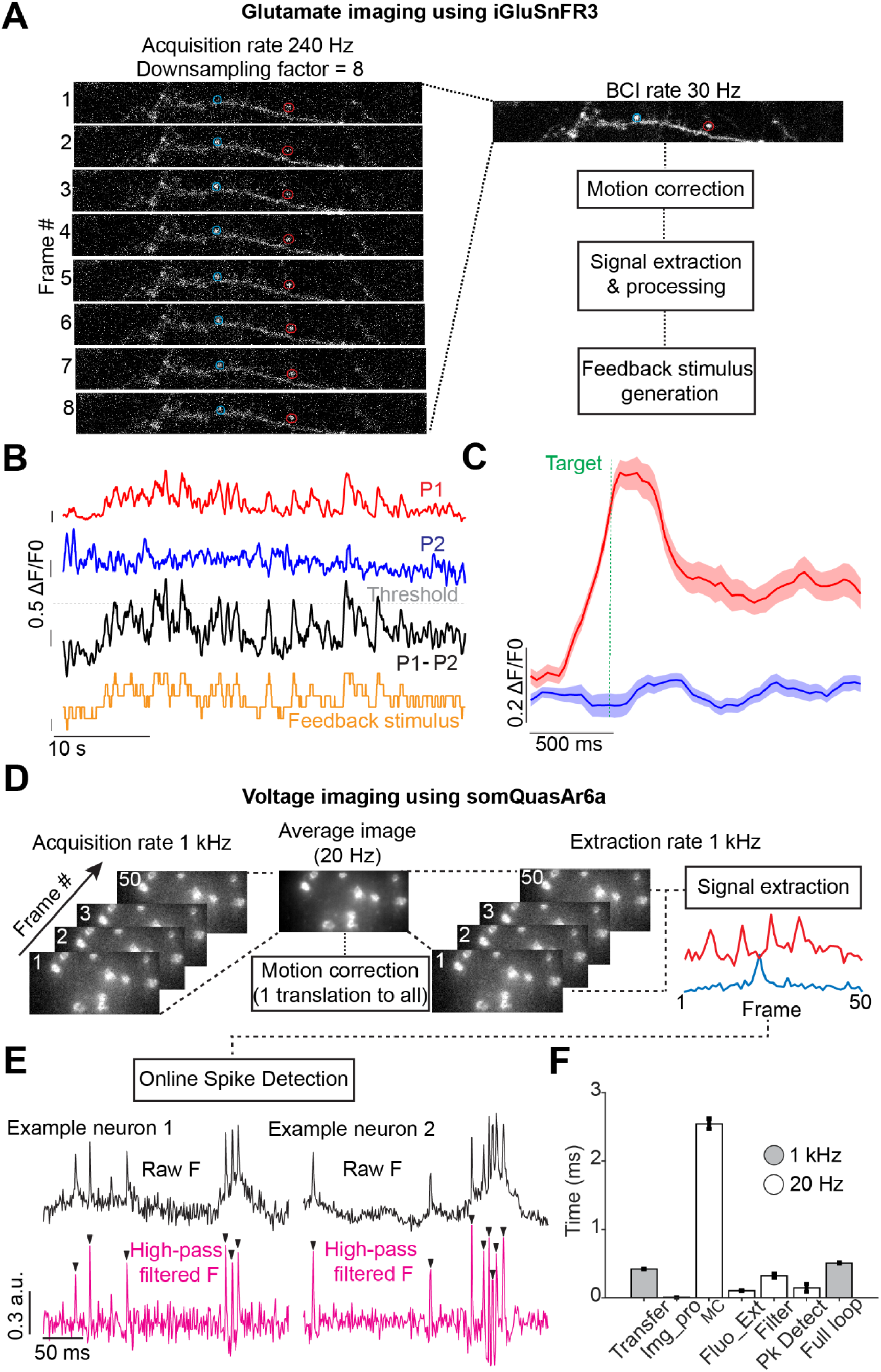
CLOOSE for BCI experiments using superfast indicators for glutamate and voltage imaging. **A**, Dendrites of a layer 2/3 pyramidal neuron labelled with iGluSnfr3. The 8 images have been acquired at 240 Hz and averaged together, effectively downsampling and at the same time low-pass filtering the images to 30 Hz. All the following steps are executed by CLOOSE at this effective imaging rate. **B**, Raw traces (at 30 Hz) for the two spines, P1 and P2, highlighted in red and blue respectively in a. In black, their subtracted signal, used as the primary signal to run the BCI. The grey dashed line indicates threshold. In orange, the stimulus feedback. **C**, Average ΔF/F0 for P1 (in red) and P2 (in blue) spines at target during a BCI session. **D,E,** A second, independent approach to perform BCI on images acquired at ultrafast speeds. D shows somQuasar6-labelled neurons under one-photon microscopy and the data processing pipeline. Images are acquired at 1 kHz and downsampled by a factor of 50 down to 20 Hz, the effective frequency at which the BCI is run. First, CLOOSE averages 50 images together and uses the average image for motion correction. Once the x and y displacement for the average image is estimated, the translation is retroactively applied to all the 50 images and the fluorescent traces for all the ROIs causally controlling the BCI is extracted for all the 50 frames. In e, this signal is first high-pass filtered (at frequencies that are user-defined, 50 Hz cutoff in this example) and after that, a peak detection algorithm is applied to detect individual spikes. Detected spikes are used to estimate an instantaneous firing rate which is mapped onto a feedback stimulus during the closed-loop part of the BCI. **F**, Time latencies for various processing steps of the voltage imaging data. In grey, the steps processed at 1 kHz (includes image processing (Img_pro)). In white, the steps processed at 20 Hz. The image (96 x 200 pixels) is transferred internally between two instances of MATLAB at full quality (uint16). Motion correction is performed using a single quadrant as large as the full image (96 x 200 pixels). Full loop is the average it takes CLOOSE to fully process a single frame, including transferring (0.4 ms). This is less than half the acquisition rate (1 ms).

### CLOOSE for BCI experiments using superfast indicators for glutamate and voltage imaging

Data acquisition from ultrafast indicators (for imaging at 100Hz or above) poses significant challenges to BCI experiments. First, high acquisition rates mean that online image processing and analysis must be completed at latencies shorter than the latencies between the acquisition of any two consecutive frames. Second, the subject’s perceptual threshold for sensory stimuli can be significantly lower than the acquisition rates of the imaging data^42^, posing the question of how to downsample neuronal activity appropriately to transform it into a perceptible sensory stimulus. On top of this, super-fast indicators might generate response kernels whose half-width is too narrow to be satisfactorily captured with standard fluorescence metrics (i.e., ΔF/F_0_) and might require additional signal processing such as spike detection – creating an additional computational bottleneck.

To address these challenges, we endowed CLOOSE with multiple complementary approaches for data downsampling and signal processing. In its most basic form, CLOOSE downsamples incoming data simply by averaging (pixel by pixel mean intensity) every n^th^ frame and processing the data at 1/n of the original acquisition speed (Figure 5a). This approach is particularly advantageous for accurate motion correction of sparsely-labelled, low signal-to-noise ratio (SNR) data which will be averaged before image registration, effectively increasing the SNR of each re-aligned frame and improving the accuracy of the registration. For example, for every 8 frames collected at 240 Hz, an average of these 8 frames will be used for both motion correction and average signal extraction (Figure 5a-c).

While this approach may work well for indicators with activity kernels of a few tens of milliseconds (e.g. iGluSnfr3^43^), it might not be as effective for indicators with even faster activity kernels, such as voltage indicators. This is because very fast fluorescence changes (i.e., a spike) might be averaged out preventing an accurate estimation of firing rates. As a consequence, faster indicators require both the raw fluorescence signal to not be downsampled and a different approach to fluorescent analysis compared to ΔF/F0. An additional challenge arises from performing computationally-intensive motion correction at frequencies compatible with voltage imaging. To overcome this, we designed CLOOSE to perform image registration and signal extraction at different frequencies. Additionally we endow it to perform online spike detection from fluorescent signals (Figure 5d, e). Specifically, users have control over a motion-correction downsampling parameter which effectively determines the frequency of motion correction. CLOOSE will perform image registration at a user-specified frequency using an average image of d_mc_ frames, where d_mc_ equals the downsampling factor. Once the x and y image translation is estimated, the translation will be applied to all the d_mc_ frames used to generate the average image and a fluorescent signal will be extracted for each frame (Figure 5d). Using this approach, users can perform accurate motion correction, maintaining the advantage provided by estimating the translation on an average image in low SNR conditions, while preserving a fluorescent signal with single frame resolution. In figure 5d-f we used this approach to perform motion correction at 20Hz, while imaging and performing spike detection at 1 kHz with an average loop latency of 0.4 ms. While averaging before motion correction leaves motion within an averaged batch to be unaccounted for, CLOOSE can perform accurate motion correction at frequencies between 300 and 500 Hz (Figure 1f and 5f). This is well above the frequency of brain motion, making residual movements negligible for most (if not all) practical applications. Once the fluorescent signal is extracted, CLOOSE will perform online spike detection on the raw fluorescence traces. This is achieved by first applying a high-pass filter to the fluorescent signals (the frequency cutoff is user-defined) and then applying a peak detection algorithm to the filtered signal to detect an instantaneous firing rate of each individual ROI causally controlling the BCI (Figure 5e). The versality of CLOOSE allows users to perform closed-loop experiments using a wide range of imaging systems and indicators for experimentalists with minimal coding skills at unprecedented speeds.

### A Graphical User Interface for easy experimental parametrization

To facilitate the user experience, CLOOSE can be parameterized with an easy-to-use graphical user interface (GUI). The GUI’s display panels endow the experimenter with visual intuition of the software’s hidden processes, including the plotting of fluorescent traces, the feedback received by the experimental model, and online image registration (Figure 6). With this interactive tool, researchers can swiftly initiate data acquisition with optimal workflow efficiency, thereby reducing the duration of experimental sessions: an advantage for both the experimenter and the animal model. The GUI’s parameters are comprehensively described in the documentation page associated with CLOOSE.

**Figure 6:**
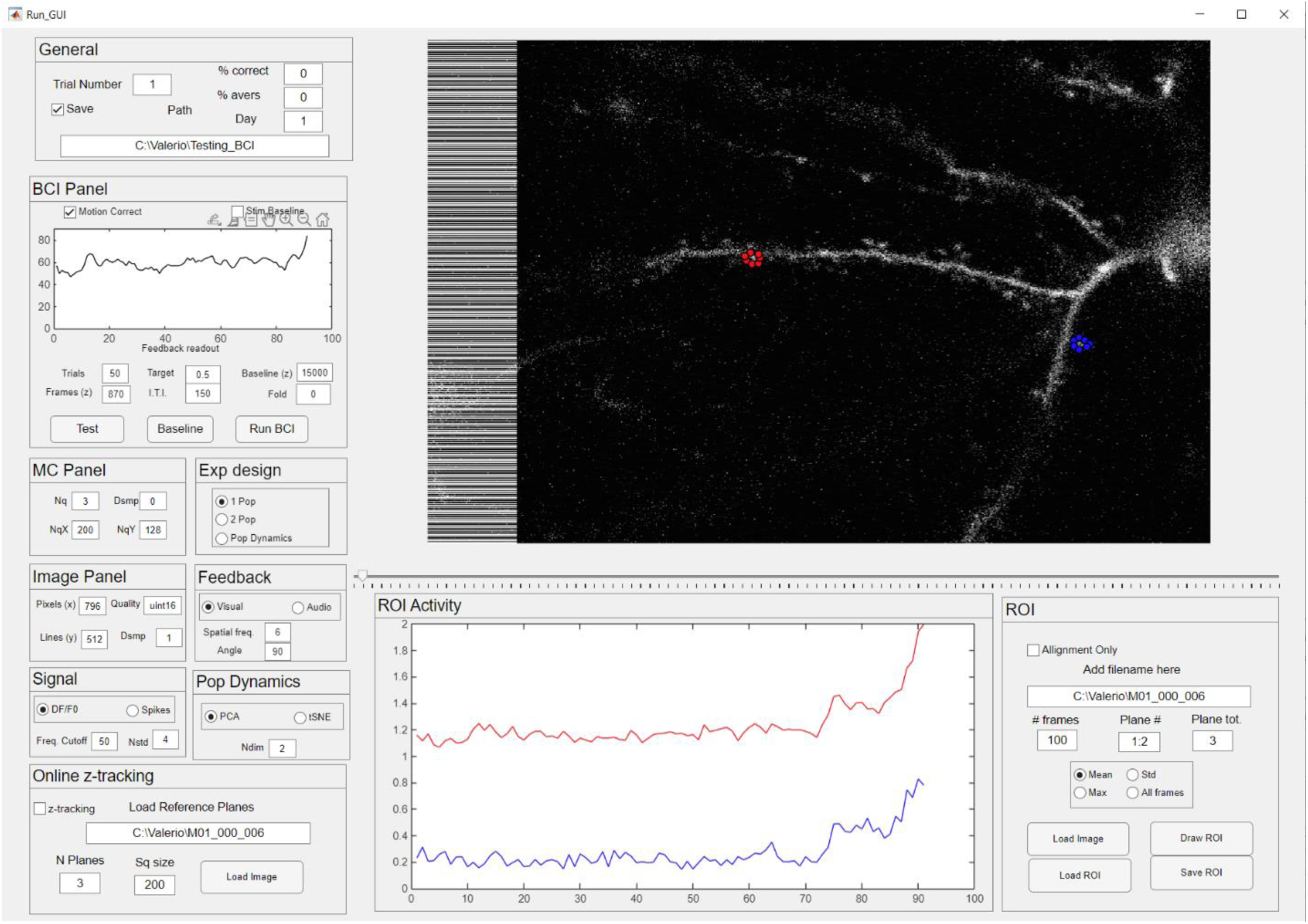
A Graphical User Interface for easy experimental parametrization. A, screenshot of the CLOOSE GUI. By interfacing with this GUI, experimenters can easily customize their experiments. These customizations include: which task design they decide to run; how many imaging planes will be used for running the BCI; what kind of signal to use to run the experiment (i.e. ΔF/F0 vs spike inference); the downsampling rate for both imaging and motion correction; the dimensions and quality of raw data, and others described in our wiki. Users can also use the CLOOSE GUI for ROI drawing on their reference image as well as to visualize their incoming data, the fluorescent traces of their ROIs and the feedback received by their experimental model.

## Discussion

We report the development of CLOOSE, a versatile, open-source MATLAB-based platform designed to standardize and democratize closed-loop optical experiments in neuroscience. CLOOSE addresses several longstanding challenges inherent to closed-loop systems—from the steep learning curves and technical complexities of synchronizing diverse hardware components to the limitations imposed by proprietary software on code accessibility and customization. Our results demonstrate that CLOOSE facilitates the online processing and analysis of high-speed imaging data and also supports operating a wide range of experimental paradigms, from single neuron to population dynamics and feedback paradigms. One of the most significant advantages of CLOOSE is its modular architecture and compatibility with virtually any imaging acquisition system via a standardized TCP/IP connection. This design allows researchers to integrate CLOOSE seamlessly with any existing frame-based image acquisition system including one- and two-photon microscopes, line and raster scanning as well as fast volume imaging using a Piezo or an ETL. CLOOSE provides the flexibility to customize key parameters for motion correction, fluorescence extraction, and signal processing, allowing the experimenter to easily tailor the software’s parameters to their experimental needs. This flexibility encourages the adoption of closed-loop methodologies by significantly lowering the technical barriers for researchers with varying levels of expertise in software engineering and signal processing.

The performance benchmarks we present demonstrate CLOOSE’s ability to handle large-scale neuronal datasets with minimal latency. With motion correction executed at subpixel precision and complete processing loops occurring at sub-millisecond latencies, the platform ensures that closed-loop feedback is delivered in a timely manner. Additionally, the capability to run different steps of the online analysis at different frequencies further allows researchers to tailor the system to the dynamic properties of the indicators used for reading out neuronal activity.

Beyond its technical merits, CLOOSE is a major step toward the widespread application of sophisticated closed-loop technology in neuroscience. By offering detailed documentation, user-friendly interfaces, and open-source code, we aim to foster a collaborative environment that accelerates the adoption and customization of closed-loop systems. Such standardization is expected to enhance data reproducibility and facilitate multi-site collaborations, ultimately contributing to more robust and comparable findings across studies. The ability to precisely control external devices based on neuronal activity not only augments experimental precision but also opens new avenues for investigating learning, synaptic plasticity, and the neuronal underpinnings of behavior in health and disease.

Despite its many advantages, CLOOSE does have limitations that warrant discussion. While its source code is publicly accessible, CLOOSE is developed in MATLAB, a proprietary software requiring a license. However, given that CLOOSE is primarily designed for use with functional biological imaging systems, its target users are research labs likely to already have access to MATLAB through their institutions. Future implementations of CLOOSE may also be developed in non-proprietary languages such as Python. Secondly, despite its ease of use, unlocking CLOOSE’s full potential still requires the user to write some (minimal) code to stream data from the acquisition system to CLOOSE. To cross this barrier, we provide example snippets of code in MATLAB and Python (Supplementary Figure 1) which the user can adopt to synchronize CLOOSE with their acquisition system.

Overall, CLOOSE represents a significant advance in the field of optical BCIs. By combining high-speed and precise online analysis with a user-friendly and modular design, CLOOSE will facilitate the widespread adoption of closed-loop functional analysis for neurofeedback studies and for the online readout and manipulation of neuronal activity.

## Method

### Data transfer to CLOOSE using a TCP/IP connection

CLOOSE is written using App Designer (in MATLAB 2024b) for building its GUI and depends on standard MATLAB packages including the Image Processing Toolbox, the Statistics and Machine Learning Toolbox and the Instrument Control Toolbox. For visual feedback, users need to have the Psychtoolbox installed too^39^. Data are streamed to CLOOSE from any image acquisition device via a TCP/IP connection. The code for this communication is already included in the CLOOSE main software while example code to mimic an acquisition device is provided as a snapshot in Supplementary Figure 1, and as actual code in the main CLOOSE repository on GitHub. This code is provided both in Matlab and Python. .bin or .tiff data can be transferred as a column vector of pixel intensities in either uint8 (1 byte) or uint16 (2 bytes) data format to control image quality and speed transfer. To setup the connection between two different PCs, both machines should connect to the same network either via ethernet or WiFi (slower). Users can use their PCs IPv4 addresses and the same port number which can be set to anything available (e.g. 5000 or 8080). For communication between two programs running on the same PC, users can use the internal IP address 127.0.0.1 (localhost). Users can find out their local IP address by typing ipconfig/all on their command window. Upon receiving a .bin or .tiff column vector, CLOOSE will reconstruct a 2D image using user-defined parameters (lines, columns and data format) specified by the user via the GUI. For each user-defined ROI, CLOOSE will extract a mean fluorescence value from all the individual pixels belonging to a given mask. ROI masks can either be drawn before the beginning of an experiment following the instructions in the GitHub wiki, or uploaded from a previous session.

### Online motion correction

CLOOSE motion correction algorithm is adapted from Sicairos *et al.* (2008)^37^. Every new image is compared to a reference image using an approach based on nonlinear optimization and discrete Fourier transforms that allow accurate subpixel registration of two images with large upsampling factors. In brief, the input images are represented in the Fourier domain:

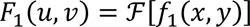

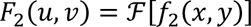

where f_1_(x, y) is the reference image, f_2_(x, y) is the image to be registered, and *Ƒ* denotes the Fourier Transform. The cross-power spectrum is computed as

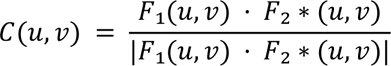

Where 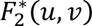 is the complex conjugate of 𝐹_2_(𝑢, 𝑣). The denominator ensures normalization.

Taking the inverse Fourier transform of C(u, v), we obtain the cross-correlation function in the spatial domain:

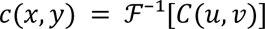

The peak of c(x, y) gives an estimate of the translation between the images. For subpixel accuracy, the algorithm performs an upsampling step using a Discrete Fourier Transform (DFT) computed via matrix multiplication:

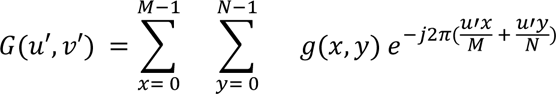

Where G(u′, v′) is the upsampled DFT, g(x, y) is the original image and (u′, v′) denotes the finer frequency grid. This refines the shift estimate to within 1/𝑢𝑠𝑓𝑎𝑐 of a pixel where usfac is the upsampling factor. The registration error is computed as:

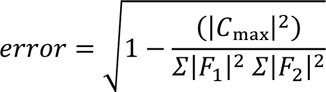

The global phase difference is given by:

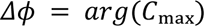

where 𝐶ₘₐₓ is the peak value of the cross-correlation. Once the shifts (𝛥𝑥, 𝛥𝑦) are found, the registered version of 𝐹_2_(𝑢, 𝑣) is computed as:

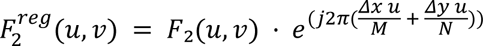

which compensates for the computed shift. This estimate is computed for each subframe. The number and dimensions (lines x rows) of each subframe is user-defined and depends on the computational speed required by the user according to their specific experimental design.

### Animals

All experiments were compliant with guidance and regulations from the NIH and the Massachusetts Institute of Technology Committee on Animal Care. Male and female C57BL/6 and Rbp4-Cre heterozygous mice were maintained on a 12-hour light/dark cycle in a temperature- and humidity-controlled room with ad libitum food access and were used for experiments at 8-15 weeks of age.

### Cortical Surgeries for GCaMP7f and iGluSnfr3 imaging

Mice were anaesthetized using 4% isoflurane and subsequently maintained at 1-2% isoflurane throughout surgery. Body temperature was maintained at physiological levels using a closed-loop heating pad. Additional heating via a heating pad was provided until full post-surgical recovery. To protect eyes from dryness, eye cream (Bepanthen, Bayer) was applied throughout the length of the procedure. At the beginning of the procedure, animals were injected with Dexamethasone (4mg/kg), Carprophen (5mg/kg) and Buprenorphine (slow release, 0.5mg/kg) subcutaneously. Hair on the scalp was removing by shaving and using hair-removal cream. The scalp was cleaned using alternating passes of iodine solution and ethanol, and the skull was exposed. For in vivo imaging, a 3mm-wide craniotomy was performed. For layer 5 labelling with GCaMP7f in Rbp4-Cre^+/-^ mice 100 nl of AVV1-syn-FLEX-jGCaMP7f-WPRE (Addgene, catalog # 104492-AAV1, 2-5x10^12^ vg/ml concentration after a 1:10 dilution from the original concentration) was injected at 400 μm from the surface of the brain at 3-4 different sites. For layer 2/3 labeling with iGluSnfr3, in C57BL/6 animal we co-injected 100 nl of AAV1.CamKII 0.4.Cre.SV40 (Addgene, catalog # 105558, 2-5x10^9^ vg/ml concentration after a 1:2000 dilution from the original concentration) and AAV.hSyn.FLEX.iGluSnFR3.v857.GPI.codonopt (Addgene, catalog # 175181, 2-5x10^12^ vg/ml concentration after a 1:10 dilution from the original concentration) at 250 μm from the surface of the brain at 3-4 injection sites. Injection speed was set at 30 nl/m, using a Stoelting motorized injector. To penetrate the dura avoiding dimpling, glass pipettes for injections were beveled beforehand. All injections sites were in the left hemisphere of the Retrosplenial cortex (2.5 mm caudal of bregma). During all surgeries the dura was left intact. Cranial windows consisted of one 3mm coverslip stacked on top of a 3 mm glass ring attached to a larger 5 mm glass ring which was subsequently fixed to the skull using cyanoacrylate glue and dental cement. A custom metal headplate was implanted in order to perform imaging under head-fixed conditions. At the end of the procedure, a single dose of 25mL/kg of Ringer’s solution was injected subcutaneously to rehydrate the animal. Animals were monitored closely for 4 days post-surgery. Recordings were started 4-6 weeks after surgery.

### CA1 Surgeries for voltage imaging

The procedures for surgeries and imaging in CA1 were based on the protocol from Dombeck et al. 2010^44^. For voltage imaging experiments viral mixtures consisted of AAV2/9 hSyn-fDIO-somQuasAr6a-EGFP-P2A-sombC1C2 (final concentration ∼2×10^12^ GC/mL) mixed with AAV2/8 CaMKIIα-Flpo (final concentration ∼0.5 – 1×10^10^ GC/mL). Virus were injected in the right hippocampal CA1 (250 – 300 nl, 45 – 60 nL/min, AP: -2.1 mm, ML: 1.9 mm, DV: -1.45 mm). Virus was injected and then a 3 mm round craniotomy (centered at AP: -2 mm, ML: 2 mm) was opened using a biopsy punch (Miltex). The dura was then gently removed, and the overlying cortex was aspirated using a blunt aspiration needle under constant irrigation with cold PBS. The center region of the external capsule was removed to expose hippocampal CA1. A cannula window was prepared prior to the surgery and comprised a 1.5 mm long stainless-steel tube (3 mm outer diameter, MicroGroup) and 3 mm round #1 cover glass (Harvard Apparatus) glued together with UV curable adhesive (Norland Products, NOA 81). Once bleeding stopped, the cannula was then lowered onto the CA1 surface until the window touched the tissue. The remaining outer surface of the cannula was sealed to the exposed skull with cyanoacrylate adhesive and dental cement that was dyed black using black ink (C&B metabond, Parkell, No. 242-3200). Finally, a stainless-steel head plate was fixed onto the exposed skull. Animals were placed on a warming blanket to recover.

### Two photon imaging with GCaMP7f and iGluSfr3

Imaging was performed using a Neurolabware two-photon microscope equipped with GaAsP photomultiplier tubes. Imaging was performed at 980 nm using an ultrafast pulsed laser (Spectra-Physics, Insight DeepSee) coupled to a 4x pulse splitter to reduce photodamage and bleaching. For excitation and photon collection we used a 16x Nikon objective with 0.8 numerical aperture for GCaMP7f imaging and a x25 Olympus objective with 1.05 numerical aperture for iGluSnfr3. For GCaMP7f imaging, we performed bidirectional scanning (512-796 pixels) semi-simultaneously in two separate planes using an electrically-tunable lens at 30.92 Hz (15.46 Hz for each plane). For iGluSnfr3 imaging, we performed bidirectional scanning (85-796 pixels) at 247.36 Hz. Laser intensity was independently optimized for each imaging session and for each plane using an electro-optical modulator. A custom light shield was attached to the headplate in order to avoid external light contamination. Animals were habituated to human handling for 5-10 minutes every day and to head-fixation for 15 minutes a day for at least three days preceding imaging. For imaging of layer 5 neurons, to maximize the number of units and to reduce signal contamination at the same time, we imaged the trunk of layer 5 pyramidal neurons at two different planes: proximal to the soma and right below the nexus (tuft bifurcation point).

#### One photon voltage imaging with SomQuasar6a

The optical system for voltage imaging consisted of a red laser (λ = 639 nm) path for holographic targeted illumination and a two-photon path for structural imaging, as described in Fan *et al*. 2023 ^45^. In brief, a red laser (CNI Inc., MLL-FN-639, λ = 639 nm, 1000 mW single transverse mode) was attenuated with a half-wave plate and polarizing beam splitter, expanded to a collimated beam of ∼42 mm diameter, then projected onto the surface of a reflection-mode liquid crystal spatial light modulator (BNS) as with the macroSLM with a resolution of 1536×1536 pixels. Polarization of the beam was set with a zero-order half-wave plate. Zero-order diffraction was blocked by a custom anti-pinhole comprised of two magnetic beads (K&J Magnetics, D101-N52) on each side of a glass slide (VWR, Menzel Glaser, 630–2129), placed in a plane conjugate to the sample image plane. The SLM was re-imaged onto the back-focal plane of the objective via a series of relay optics and the Bruker two-photon microscope. Objective lenses were a 25× water immersion objective with NA 1.05 (Olympus, XLPLN25XWMP2), and a 25× water immersion objective with numerical aperture 1.00 (Leica, HC IRAPO L 25x/1.00 W motCORR). A mechanical shutter blocked the red laser between data acquisitions and a series of OD filters were placed after the red laser for modulating intensity. The SLM device was controlled by custom software. A user specified area for the SLM to target by drawing on a 2P fluorescence image. The SLM phased pattern was calculated using the Gerchberg-Saxton algorithm. Red laser intensity was ∼ 2.5 – 3 mW per cell for *in vivo* imaging.

### Different brain computer interface tasks designs

Neurofeedback brain-computer interface learning paradigms are tasks in which mice are trained to modulate their own brain activity in order to obtain rewards. Mice receive feedback about their performance through a stimulus driven by brain activity. CLOOSE assumes that the experimenter will record a baseline (of a user-defined length – either everyday (recommended) or on day 1 of training) before the closed-loop part of the experiment to estimate target. CLOOSE allows the experimenter to run either a ‘dark’ baseline recording in which no stimuli are presented, or a ‘feedback’ baseline in which the same stimuli presented during the closed-loop task will be presented in random order. The baseline recording is necessary to estimate baseline activity which will be used to determine the neuronal activity necessary to reach target, as well as the mapping between neuronal activity and the feedback stimulus. The mapping between neuronal activity, feedback stimulus, and target can be estimated in three different ways according to the task design.

For the one population and the two populations task design, P1 and P2 activity refers to the activity of neurons which bring the feedback stimulus closer to the target, and vice versa, respectively. During baseline, the responses of P1 and P2 neurons is recorded for a period of set by the experimenter. In the one population design, P2 represents a single ROI drawn over the neuropil. For the populations task design, the P2 neurons can be either a single neuron or a population of neurons. For accurate results we recommend recording a baseline of at least 10 minutes. ΔF/F_0_ or spike rate is calculated for all P1 neurons and P2 neurons, averaged and the activity (ΔF/F_0_ or spike rates) of the two populations of neurons will be subtracted from one another. Next, CLOOSE will resample 200 trials (trial length is a user-defined parameter) from the baseline recording and iteratively search (in 0.005 ΔF/F_0_ incremental steps) for the subtracted activity value producing a user-defined baseline success rate (e.g., 50% of the trials). That value is set as the threshold value for target activity during the closed-loop phase of the BCI task. The feedback stimulus is then divided in a user-defined number of bins. The mapping between neuronal activity and stimulus feedback is linear and defined as follows: the first stimulus bin corresponds to the minimum value in the subtracted ΔF/F_0_ signal or spike rate distribution, while the target is reached at the subtracted ΔF/F_0_ or spike rate value corresponding to threshold defined as described above. Activity in between is split into equally spaced bins. The feedback stimulus is updated at each new incoming frame (or downsampled frequency, as defined by the user) as the subtracted ΔF/F_0_ signal or spike rate averaged over the last three frames.

Similarly, for the network dynamics BCI design, during baseline the responses of P_nd_ neurons is recorded for a period of time which is going to be set by the experimenter. For accurate results we recommend recording a baseline of at least 10 minutes. ΔF/F_0_ or spike rate is calculated for all P_nd_ neurons for the entire duration of the baseline and z-scored in a neuron-by-neuron manner. Z-scored activity will then be collapsed onto an n-dimensional space where the dimensionality reduction method can be either PCA or tSNE and n is user-defined for PCA equal to 2 for tSNE. Next, CLOOSE will resample 200 trials (trial length is a user-defined parameter) from the baseline recording and use the highest value in the first dimension of the reduced space to find the n-dimensional point (P_ref_) where neuronal activity would reach target (P_targ_). From now on, P_ref_ will be the point in relation to which all Euclidean distances will be estimated. To find target activity, CLOOSE will first find the furthest point in the dimensionally-reduced space from Pref and set this distance (P_dist_) as the first stimulus bin. The edges of the target bin are found by decreasing P_dist_ in decrements of 100^th^ of its value, until a value (P_targ_) is found for which a user-defined fraction of the resampled trials (e.g. 50%) is crossed. Activity falling at distances from Pref between P_targ_ and P_dist_ is split into equally spaced bins and mapped to a feedback stimulus.

### Volumetric BCI

For volumetric BCI, the user will first acquire a reference image from each plane using an ETL or a Piezoelectric motor. Next, CLOOSE will prompt the user to draw ROIs on each of the recorded planes. The user may decide to draw ROIs on all planes, or just on a subset of them. During the acquisition phase, the reference image from the first plane will be used to motion correct all the other planes. CLOOSE will extract fluorescence from the ROIs in the planes where the user drew ROIs and ignore the planes with no ROIs.

### Online z-tracking

For online z-tracking, users can either acquire or upload reference volume images from a user-defined number of planes. During acquisition the incoming frame will be registered in x and y against the reference image as described above (*Online motion correction* method section). At the beginning of the recording, an estimated x and y shift, will be applied to the entire volume to re-align the z-stack with the incoming data. The x and y shift will be calculated by registering the z-stack frame with the lowest MSE from the reference image, against the reference image itself. Once re-aligned, the MSE between the new frame and all reference frames in the z-stack will be estimated using a quadrant of size S_q_ (user-defined). During acquisition, users have the option to live-plot the inferred z-shift for online tracking. For the data shown in Figure 4, we generated ground-truth data by imaging static pollen at 21 different depths using an ETL. We then compared each of these images, acquired at different depths, against the 21 reference images.

### Benchmark tests

Benchmark tests were performed as follows: For testing the speed of data transfer illustrated in Figure 1b-d, first we generated a synthetic image dataset using MATLAB of different sizes (512 x 796 and 256 x 796, full and half image size, respectively) of different image qualities (uint8 and uint16). Next, we tested the speed at which we transferred these images: for images transferred within the same PC, we transferred the images across two instances of MATLAB, via an internal connection (localhost). For images transferred across the network between 2 different PCs, we transferred the images across two instances of MATLAB, each of which opened on a separate machine, via an internal network using an ethernet connection. In both instances, we transferred 10000 frames. Data were transmitted between two PCs using a direct Gigabit Ethernet via a Netgear ProSAFE GS105 Gigabit switch, which ensured stable, high-throughput communication connection and the TCP/IP protocol via MATLAB’s tcpip objects. PC1 (the acquisition device) had the following specifications: Intel Xeon W-2223, 128 GB RAM, Windows 11. PC2 (CLOOSE device) had the following specifications: Intel(R) Core (TM) i7, 32 GB RAM, Windows 10). To do so, we used a custom MATLAB script. Each frame was sent as a 1D uint16 vector. The transfer speed was estimated using an internal clock on the receiving PC. Additionally, we tested the speed of different processing stages including image processing, motion correction, fluorescence extraction, plotting and stimulus presentation. These steps where tested internally within CLOOSE, by setting various clocks (tic and toc functions in MATLAB) at the beginning and end of the specific part of the loop tested. The speed of these steps was quantified on an Intel(R) Core(TM) i7-10700k with a CPU operating at 3.8 GHz with 32 Gb of RAM (PC2, as above). The accuracy of motion correction was estimated by comparing the performance of online rigid motion correction using CLOOSE against the offline output of Suite2P for rigid motion correction. Finally, for estimating the performance of CLOOSE on z-tracking in Figure 4, we generated ground-truth data by imaging static pollen at 21 different depths, separated by 1 μm, using an ETL (known depth). We then compared each of these images, against 21 reference images, also acquired using an ETL at the corresponding 21 planes. An error was defined as the distance between the known ground truth, and the estimated z-plane.

### BCI downsampling

Downsampling is necessary in order to produce sensory feedback at rates which are within the mouse perceptible range. Users can downsample two independent variables: image processing and motion correction rate. Image downsampling consists in averaging multiple frames together and then run all thew following analysis on an average image. This option is useful when running BCI experiments using indicators which require high sampling rates (>50 Hz) and with low SNR (e.g., iGluSnfr3). The motion correction downsampling on the other hand, averages a use-defined set of images for motion correction only. Images will be first averaged together. Next, the average image will be used for estimating the x and y displacement against a reference image. This translation will be applied to all the images that were averaged together and the fluorescence signal will be extracted in a frame-by-frame manner. This approach heavily reduces the computational load on CLOOSE as motion correction will be applied on only a subset of loop iterations and it is therefore generally convenient to apply when imaging at high acquisition speeds.

## Acknowledgements

We thank Gabriela Rasch, Zhaoran Zang and Courtney Yaeger for comments on the manuscript. M.T.H. was supported by the NIH (RO1NS106031, R01NS113079 and R01MH135141). V.F was supported by the Y. Eva Tan Molecular Therapeutics Postdoctoral Fellowship.

## Data and code availability

All analysis, code, and data are available at https://github.com/harnett/CLOOSE. CLOOSE code and wiki are open-source and they will be deposited in full in its GitHub repository under a GNU GPL-3.0 license, upon publication of this manuscript. We welcome any user to download, inspect and use the main code.

## Contributions

V.F. conceptualized and wrote the main software, habituated and performed surgery on mice, collected the iGluSnfr3 and the GCaMP7f data, conceptualized and implemented the analyses, prepared the figures, and wrote the manuscript. A.B. habituated and performed surgeries on mice, collected the iGluSnfr3 data, and helped write the manuscript. L.Z.F. collected voltage imaging data and helped write the manuscript. M.T.H supervised all aspects of the project.

## Ethics declarations

The authors declare no competing interests.

---

**Supplementary Figure 1:**
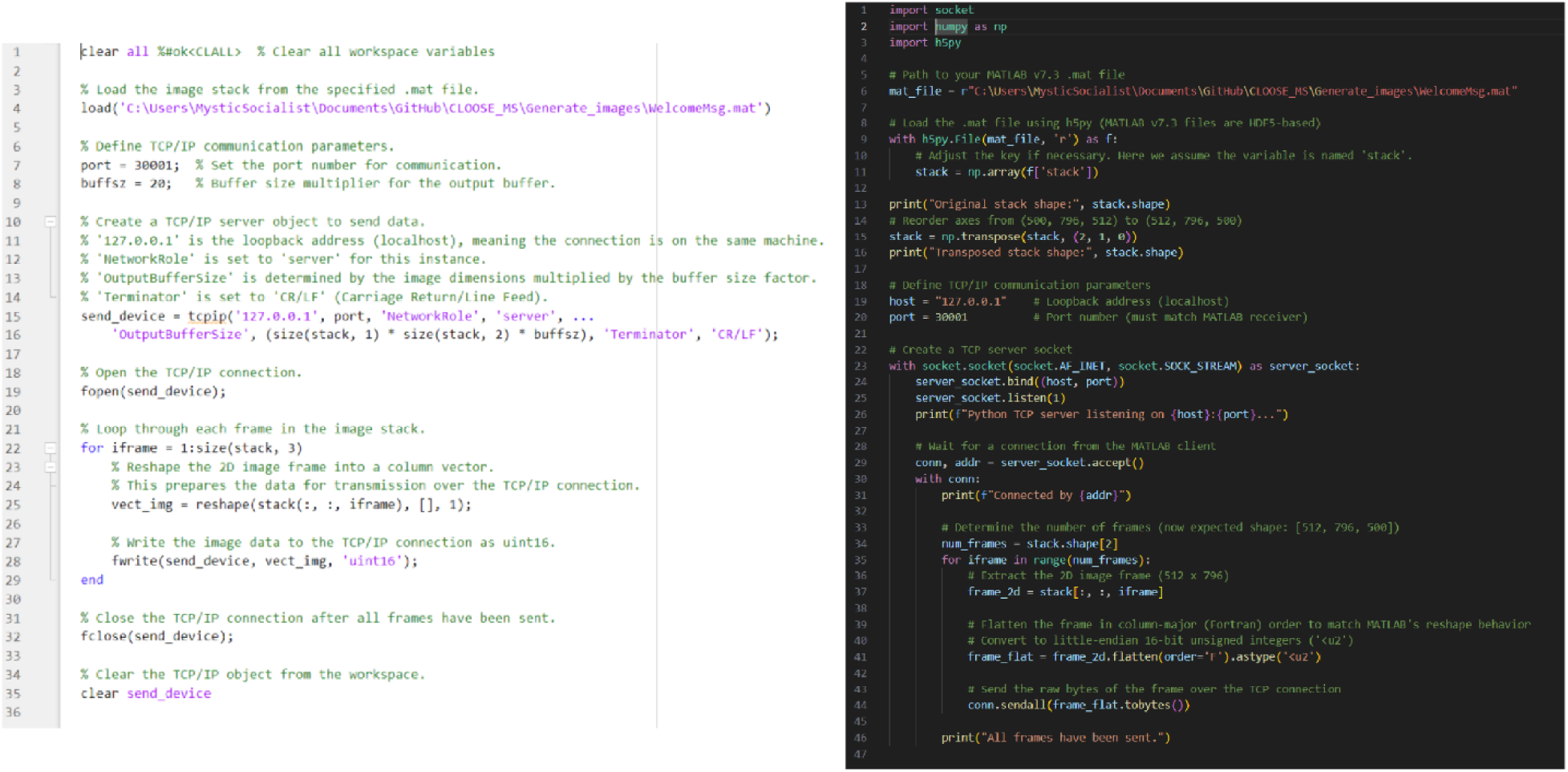
Code snippets to transfer data from an image acquisition device to CLOOSE. **A,** The actual code, in MATLAB (left) and Python (right), is provided within the CLOOSE software package. This is the bulk of the coding required to utilize CLOOSE.

